# Shift in motor-state equilibrium explains gait therapy effects of apomorphine in experimental Parkinsonism

**DOI:** 10.1101/2025.09.07.673890

**Authors:** Burce Kabaoglu, Elisa L. Garulli, Rafael De Sa, Arend Vogt, Ruben Behrsing, Matej Skrobot, Raik Paulat, Patrick Pollak, Lynn S. Guldin, Moritz Gerster, Wolf-Julian Neumann, Matthias Endres, Christoph Harms, Nikolaus Wenger

## Abstract

Gait impairments remain a major therapeutic challenge in Parkinson’s disease (PD). Apomorphine is gaining renewed clinical attention with the expanding use of pump infusion systems. Yet, the specific role of apomorphine on the neural regulation of gait has remained poorly characterized, limiting it’s targeted use for symptom-specific therapy in PD.

Here, we examined the neurobehavioral effects of apomorphine on runway locomotion in the unilateral 6-hydroxydopamine (6-OHDA) rat model. Therapeutic drug doses significantly increased total walking distance, related to reduced akinesia and prolonged gait episodes. Conversely, 3D kinematic analysis revealed reduced limb velocities under medication.

At the neural level, therapy doses selectively enhanced cortical high-gamma rhythms without substantially altering beta or low-gamma activity. Instead, beta and low-gamma oscillations were consistently suppressed during motor activity in both medication ON and OFF conditions. Neurobehavioral correlations showed that transitions into gait were facilitated by reductions in beta and low-gamma activity, whereas transitions to akinesia were primarily suppressed when high-gamma activity was elevated.

Our findings suggest that modulating cortical activity can aid ameliorating gait deficits in PD. We further propose that the complex therapy effects of apomorphine are best explained by a shift in motor-state equilibrium that is defined by the transitions of akinesia, stationary movements and gait. Together, these insights establish a mechanistic framework to guide the development of targeted gait therapies in PD.

## Introduction

Parkinson’s disease (PD) is a neurodegenerative disorder characterized by motor symptoms such as tremor, bradykinesia, and gait impairments^1^. Freezing of gait (FoG), a particularly debilitating gait disturbance, is characterized by a sudden inability to initiate or continue walking, often described by patients as feeling as though their feet are “glued” to the floor^2,3^. FoG occurrence is often associated with time periods when standard dopaminergic medication wears off, in the so called OFF periods^4^. Although medication with Levodopa or DBS can alleviate bradykinesia during gait, these therapeutic benefits often diminish over time due to treatment fluctuations and disease progression^5–7^. This unmet clinical need has urged the search for alternative gait therapies.

Apomorphine has recently gained renewed attention for treating refractory PD symptoms, including motor fluctuations, axial impairments, and gait deficits^8^. It is a dopaminergic agonist with preferential activity at D2 over D1 receptors^9^. The TOLEDO randomized controlled trial demonstrated that continuous subcutaneous apomorphine infusion significantly reduced patient-reported OFF periods^10^, contributing to the approval of subcutaneous apomorphine infusion systems in the United States in 2025^8,11^. Retrospective clinical evaluations have further indicated that apomorphine may reduce FoG occurrence^8,12^, although prospective confirmation is still lacking. Overall, the specific mechanisms by which apomorphine influences the neural regulation of gait remain unclear.

Excessive beta-band oscillatory activity (13–35 Hz) is a hallmark of PD^1,13,14^. Beta oscillations are associated with bradykinesia and rigidity^13^ and suppressed by dopaminergic medication or DBS^15^. Pogosyan et al. demonstrated that beta activity facilitates tonic muscle contraction while suppressing the initiation of new movements^16^. Based on these findings, Engel and Fries have proposed that beta oscillations encode the system’s tendency to maintain the current motor state - the “status quo”^17^. In contrast, gamma oscillations (65–90 Hz) facilitate movement and increase with DBS^14,18^. Technological advances in chronic DBS recording have further positioned neural oscillations as promising biomarkers for gait impairments in PD^19^. For example, FoG episodes have been linked to elevated low-beta oscillations in subthalamic nucleus (STN)^20^, reduced low-frequency coupling between motor cortex and STN^4^, and increased beta-gamma phase-amplitude coupling in motor cortex^21^. Guided by these findings, we set out to determine how beta-and gamma-frequency activity contributes to gait regulation under apomorphine therapy.

Rodent models of PD reproduce many pathological oscillatory signatures and gait impairments observed in humans^7,22–25^. Neural correlates of rodent bradykinesia emerge within the beta and low-gamma ranges (15–40 Hz)^22^, and are suppressed by dopaminergic medication or DBS^7,26^. Excessive high-gamma activity (70–110 Hz) accompanies dyskinesia onset^27–29^. Cortical oscillations also play a central role during walking in PD rodent models^7^. For example, unilateral 6-OHDA-lesioned rats exhibit elevated beta and low-gamma activity during treadmill walking compared with sham animals, reflecting abnormal motor-cortical synchronization associated with gait impairment^24^. Despite long standing research in PD rodent models, there has been no detailed characterization exists regarding the therapeutic effects of apomorphine on gait regulation and cortical oscillations^7^.

Here, we aimed to characterize the neurobehavioral effects apomorphine during runway locomotion in the unilateral 6-OHDA rat model of PD. We acquired high-resolution video, 3D hindlimb kinematics, and motor-cortex ECoG signals, enabling identification of three principal motor states during self-initiated locomotion: gait, akinesia, and stationary movements. To derive therapeutic relevant dosing, we titrated apomorphine amounts to promote forward locomotion while avoiding excessive body rotations. This experimental design resulted in three key observations, First, therapeutic levels of apomorphine increased the total walking distance on the runway by reducing akinesia, increasing transitions into gait, and prolonging overall gait duration. Secondly, apomorphine increased cortical high-gamma activity, whereas beta and low-gamma activity decreased during motor engagement but did not change with drug application alone. Third, neurobehavioral correlations revealed that reductions in beta and low-gamma selectively promoted transitions from stationary movement into gait, whereas elevation in high-gamma activity reduced transitions into akinesia. Taken together, these findings suggest that the therapeutic effects of apomorphine can be understood as a shift in motor-state equilibrium, defined by the durations and transition probabilities of akinesia, stationary movement, and gait. Within this framework, the functional contributions of beta and gamma oscillations can be interpreted dynamically as functions of the current motor state.

## Methods

### Ethics statement

All experimental procedures were approved by the local regulatory bodies (Federal Office for Health and Social Affairs, Berlin, Germany; approval number: G0206/19) and performed in compliance with the German Animal Welfare Act. The experimental procedures were performed on male Sprague-Dawley rats (n = 14, 300–325 g, Janvier, France). Animals were kept on a 12-hour light cycle with ad libitum access to food and water. The data of this study can be made available by the corresponding author upon request.

### Surgical Procedures and Lesioning

Unilateral lesioning of midbrain dopaminergic neurons was performed on 8-week-old rats to model Parkinson’s disease. During the surgery, anesthesia was maintained with 1-1.5% isoflurane in oxygen. The body temperature was regulated with a heating mat, and the breathing rate was monitored with an oximeter (MouseOx). Upon anesthesia induction, rats received a subcutaneous injection of carprofen (5 mg/kg) for pain management. They were then secured in a stereotactic frame (David Kopf Instruments). Local anesthesia was provided under the scalp with 2% lidocaine (B. Braun SE), and eye dehydration was prevented with Pan-Ophtal Gel. The surgical site was prepared by excising skin to fully expose the skull. The dorsoventral distance between bregma and lambda was adjusted to less than 0.05 mm for accurate targeting. Using the Paxinos coordinate system, the skull was marked and a craniotomy was performed. We used a 33-gauge blunt steel needle to deliver 1 μL of solution containing 0.05% ascorbic acid and 8 μg of 6-OHDA (Sigma-Aldrich) in 0.9% sterile saline. The injection was performed into the left medial forebrain bundle (MFB) at specified coordinates (AP: −2.6, ML: +1.6, DV: −8.4, DV measured from the skull surface) using a precision pump (Harvard Apparatus). The needle was left in place at the target site for 5 minutes following the infusion of the neurotoxin.

Electrocorticography (ECoG) electrode implantations were performed 2 weeks after the initial 6-OHDA lesion. The preparation and protective measures were identical with the first surgery with added steps for electrode implantation and head stage installation. The frontal, parietal, and interparietal bones were carefully cleared from connective tissue, rinsed with sterile saline, and dried with an air blower. The exposed skull surface was then treated with a UV-activated bonding agent (OptiBond All-in-One, Kerr). A surgical pen was used to mark the sites of the craniotomies. Another UV-activated dental composite (Charisma A1, Kulzer), was then put on top of the OptiBond layer to make a shallow well. The subsequent craniotomy and durotomy allowed for the stereotactic implantation of a custom-made stainless-steel screw (1,2mm of diameter) overlaying motor cortex (AP: +3.75, ML: ±2; Ground, AP: −12.5, ML: −2). The electrodes were attached with silver wires to a 32-channel connector (polarized nano connector, PZN, Omnetics). The electronic components were subsequently encapsulated in dental acrylic (Paladur, Kulzer). After the surgery, we allowed the animals to recover for three days under close observation and administration of daily pain medication (carprofen, 5 mg/kg s.c.).

### Apomorphine dosages

Apomorphine (Sigma-Aldrich) was administered s.c. at 0.025 mg/kg for gait therapy in the runway experiments. This therapeutic dose was determined at the beginning of experiments to promote forward locomotion without inducing excessive body rotations. For the confirmation of the unilateral 6-OHDA lesion, we additionally applied a one time apomorphine dosage of 0.05 mg/kg to look for unilateral body rotations in the cylinder test, at the end of the experimental timeline (compare cylinder results in supplementary figure 1). For both tasks, the specified amount of apomorphine was applied approx. 5 min before the start of the experiments.

### Behavioral runway experiments

Rats were handled daily starting approx. two weeks prior to 6-OHDA lesioning (experimental timeline see Fig 1A). The animals were then trained for three days on the runway to habituate themselves to the experimental environment. For all runway trials in this study, we placed a home cage with a second rat at the end of the runway. This allowed for a quick habituation of the animals and sufficient motivation to repeatedly engage in the runway task. For each run, rats were placed by an experimenter at the start of the runway. After completing the run across the entire runway, or after a fixed time of 2 min the trial was terminated by picking up the animal at its final position, shuttling it to the homecage and after approx. 3-5 seconds shuttling it back again to the start of the runway. Hindlimb kinematics were captured using a high-speed motion capture system (Vicon), which included 12 infrared cameras and two high definition video cameras, operating at a recording frequency of 200 Hz and 40 Hz, respectively^30^. Reflective markers were attached to key anatomical landmarks on both hindlimbs (iliac crest, hip, knee, ankle, front quadrant of the foot) and the shoulders (Fig 2). Kinematic parameters were analyzed and computed using custom-written MATLAB scripts^30^.

**Figure 1.**
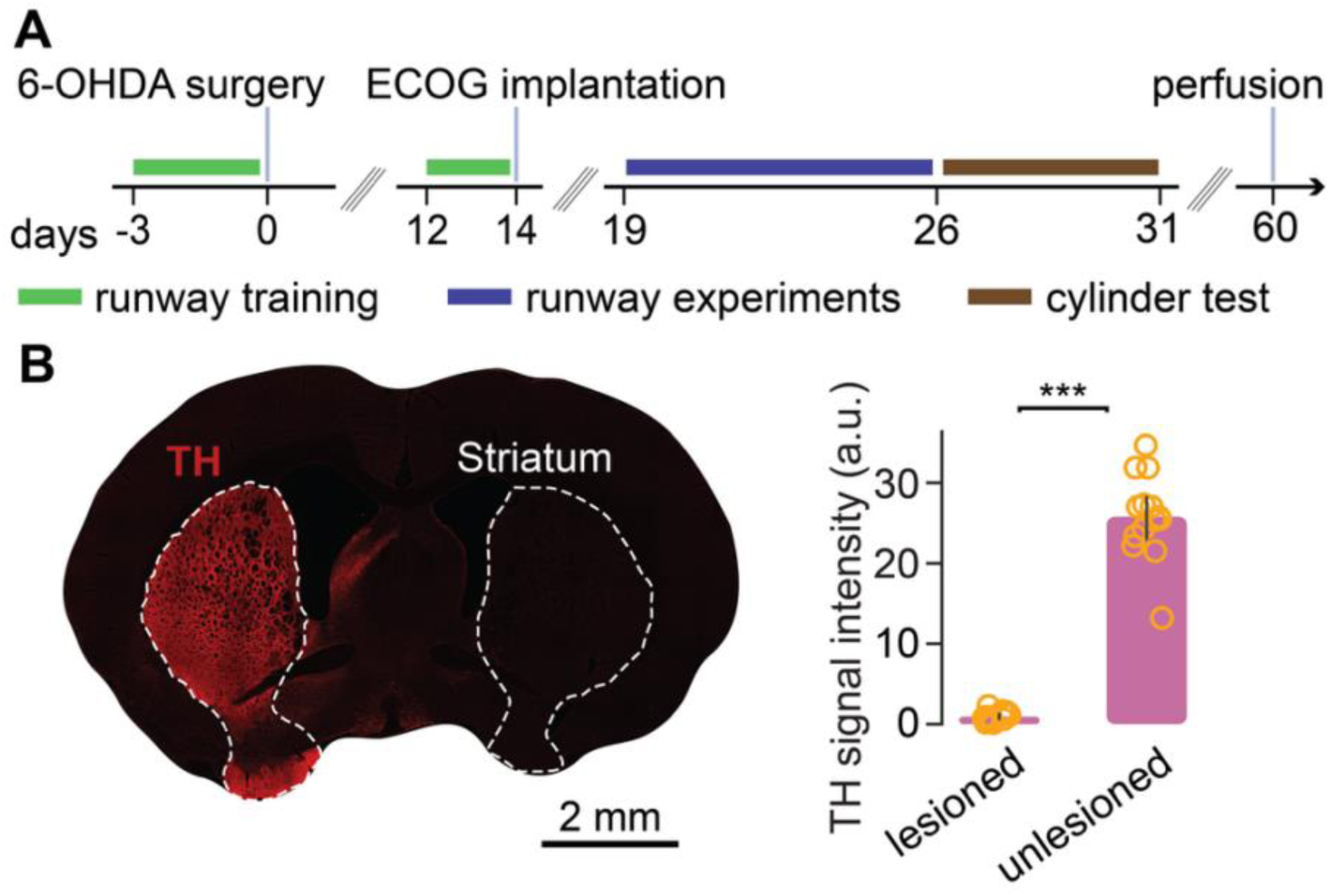
Experimental timeline and quantification of dopaminergic lesion. (**A**) Animals were habituated to the runway starting 3 days before the 6-OHDA lesion. After 12 days of recovery and 2 days of retraining, ECOG electrodes were implanted. Runway experiments were performed in the Med-OFF and apomorphine (0.025 mg/kg) conditions between day 19 and day 26 post-surgery. The 6-OHDA lesion was additionally validated using the cylinder test under a higher dose of apomorphine (0.05 mg/kg). (**B**) Histological quantification confirmed a unilateral dopaminergic lesion in the striatum. Left: representative coronal section labeled with anti–tyrosine hydroxylase (TH). Right: quantification of TH signal intensity in the lesioned versus unlesioned hemispheres (>95% reduction, n = 14 rats). Bars show mean ± SD. ***p < 0.001.

**Figure 2.**
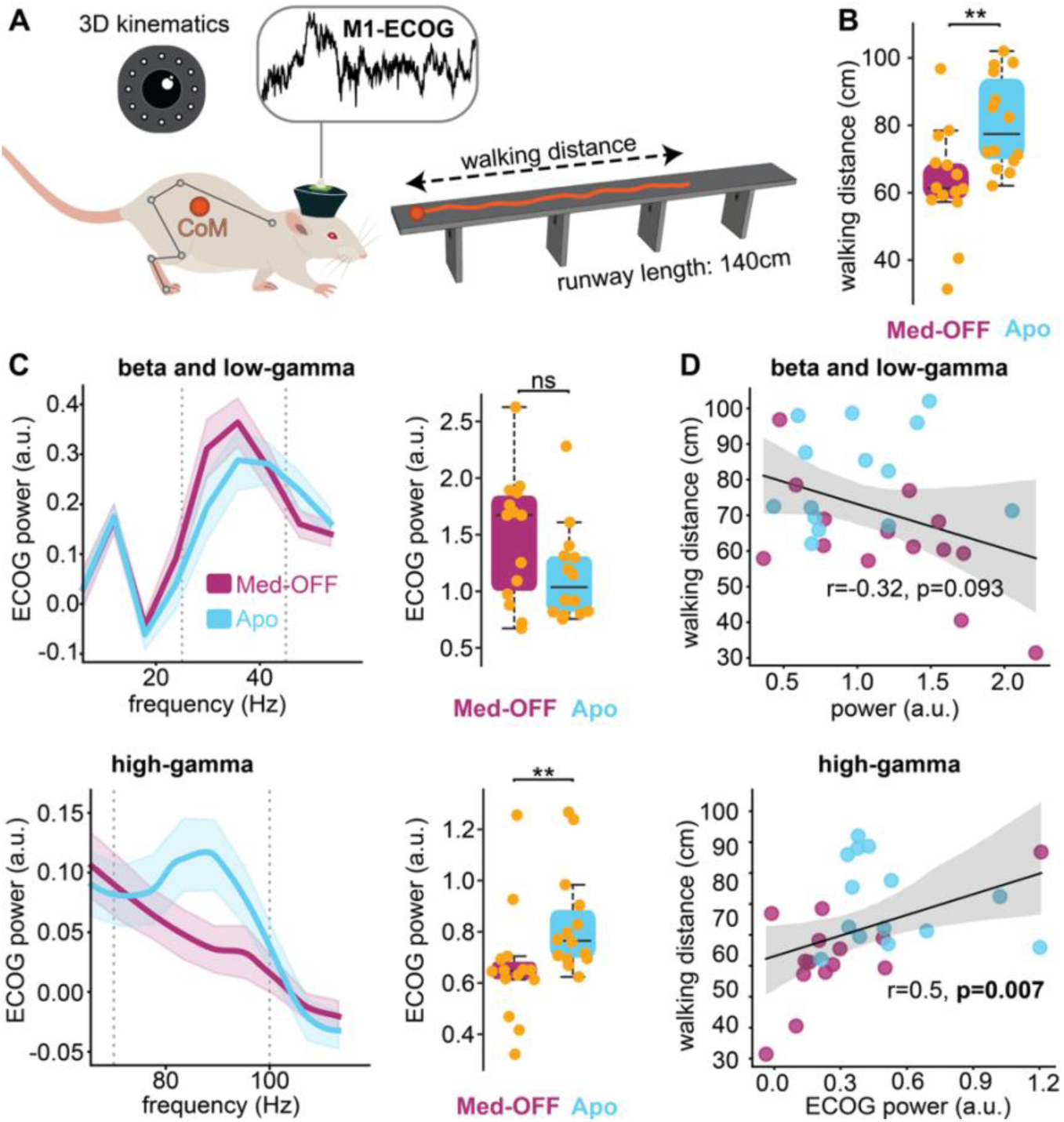
Apomorphine-induced improvements in walking distance primarily correlated with high-gamma elevations. (**A**) Experimental setup for 3D kinematic analysis and ECOG recordings on the runway. Walking distance was calculated from the trajectory of the center of mass (CoM), defined by crest, hip, and shoulder markers. (**B**) Quantification of walking distance in Med-OFF and Med-ON conditions. (**C**) Therapeutic doses of apomorphine significantly increased high-gamma activity. Gray dashed lines indicate beta–low-gamma (25–45 Hz) and high-gamma (70–100 Hz) frequency ranges. (**D**) Changes in walking distance primarily correlated with cortical high-gamma elevations. Data are presented as boxplots. Significance: ns = non-significant, **p < 0.01. Power values are reported in arbitrary units (a.u.).

Runway experiments started one week after ECOG implantation, corresponding to three weeks after 6-OHDA lesioning. In the same session, rats were recorded OFF medication and after application of a therapeutic dose of apomorphine (s.c. at 0.025 mg/kg). For each condition, rats were allowed to traverse the runway until data were collected from 15 trials. Each trial began with the animal being placed at the beginning of the runway. A trial was considered successful and terminated, if the animal traversed the runway to the other end. If the animal paused or failed to initiate walking, the trial was stopped after two minutes. Trials continued until 10 successful runs were completed or until 20 minutes had elapsed. During data analysis, videos were manually annotated in the software Nexus 2.0 (Vicon) to define the beginning and end of the different motor states: akinesia, stationary movement, and gait. These state labels were further processed with a custom pipeline built using the open-source package *neurokin* (https://github.com/ELGarulli/neurokin).

### Cylinder Test

After runway experiments finished, the effectiveness of the 6-OHDA lesion was assessed behaviorally using an apomorphine challenge within a glass cylinder. Behavior in the cylinder was documented using a video camera (ELP 1080P, Night Vision USB Camera, 30 Hz framerate). Following baseline recordings of 5 min OFF medication, we administered a dose of apomorphine at 0,05mg/kg. Five minutes after injection, the rats were returned to the glass cylinder. Trajectories of the body rotations were reconstructed using the open-source toolbox DeepLabCut^31^. We then quantified and analyzed the rotations ipsilateral and contralateral to the dopaminergic lesion using custom-written MATLAB script.

### Neural Data Acquisition

Neural data were acquired using a multicomponent system from Tucker-Davis Technologies (RZ2, TDT). The system was set to enable neural recordings at 6 kHz. Recording cables were connected to the freely behaving animals through a motorized commutator (ACO32, TDT). Neural signals were digitized locally at the head of the animal with a 32 channel digitizing headstage (Intan).

### Neural Data Analysis

Power spectral densities of M1 electrocorticograms (ECoGs) were computed using fast Fourier transform (FFT) with a Hanning window and a frequency resolution of 1 Hz. The resulting power spectra were subsequently decomposed into aperiodic and oscillatory components employing the FOOOF toolbox^32^. The FOOOF algorithm (version 1.1.0) and default settings were used for the algorithm (i.e., no custom parameter adjustments). Power spectra were parameterized across the frequency range 5 to 120 Hz. We defined specific frequency ranges for analysis: the beta and low-gamma range spanned from 25 to 45 Hz, and the high-gamma range spanned 70 to 100 Hz. The power analysis within these bands was expressed in arbitrary units (a.u.) without additional normalization.

### Histology

The extent of dopaminergic denervation was assessed post-mortem through immunohistological staining for tyrosine hydroxylase (TH). Under deep anesthesia, animals were transcardially perfused with 70 mL of 0.1 M phosphate-buffered saline containing 0.4% sodium-heparin, followed by 120 mL of 4% w/v paraformaldehyde in 0.1 M phosphate buffer. Subsequently, brains were extracted, postfixed in 4% w/v paraformaldehyde for a minimum of 24 hours, and cryoprotected sequentially in 15% and 30% sucrose solutions for 24 hours and one week, respectively. Tissues were then frozen in methylbutane (Carl Roth).

40-μm coronal cryo-sections encompassing the entire striatum from −3.24 to 2.52 relative to bregma were serially cut. Sections underwent a 30-minute preincubation in quenching solution (100 mM glycine, pH 7.4, adjusted with 2M Tris Base), followed by a 30-minute incubation in blocking solution. They were then incubated overnight at 4°C with rabbit anti-rat tyrosine hydroxylase antibody (Abcam) and anti-NeuN (neuron nucleus) antibody (Merck Millipore) diluted in blocking solution. After three 10-minute washes in PBS, the sections were incubated for 2 hours at room temperature with secondary antibodies: anti-rabbit-594 (#A-21207, Invitrogen) and anti-mouse-488 (#A-11029, Invitrogen), along with DAPI (1:10000) in blocking solution. Finally, the sections were rinsed with PBS and mounted with ImmuMount. The stained slices were imaged using a Leica DMI8 microscope, allowing detailed visualization of the stained dopaminergic neurons and nuclei.

The optical density (OD) of TH+ fibers in the striatum was measured using NIH ImageJ (version 2.9.0). OD values were calculated by subtracting the background signal (measured in the unstained corpus callosum) from both the lesioned and unlesioned striatal measurements. Three striatal sections per animal were analyzed.

### Statistics

Video raters of motor states were blinded to group allocations to minimize bias. The statistical group planning and p-values are of exploratory nature. Statistical quantification of two-factors (drug and motor state) were performed using the nonparametric Scheirer-Ray-Hare test (SHR-test). All, pairwise group comparisons were performed with the nonparametric Wilcoxon signed rank test. Correlational analysis was performed with the Spearman test. In the figures, data are plotted as individual points and boxplots reporting median and quartiles. Power spectral densities report mean ± standard deviations.

## Results

### Apomorphine improves walking distance and enhances cortical high-gamma oscillations

We first evaluated the effect of apomorphine on walking distance during runway locomotion in the unilateral 6-OHDA rat model of PD. An overview of the experimental timeline is shown in Figure 1A. Histological verification confirmed a robust loss (>95%) of TH-positive innervation in the striatum of the lesioned hemisphere (Fig. 1B). On the runway, rats received a therapeutic apomorphine dose of 0.025 mg/kg, selected to modulate network activity while promoting straight-line walking (Fig. 2). Under this dosing condition, PD rats traveled significantly farther than during medication-OFF sessions (80 cm ± 13.33 SD vs. 60 cm ± 15.20 SD, p < 0.01; Fig. 2B).

We next examined whether changes in walking distance were associated with alterations in cortical oscillations across the full duration of each runway trial. In the medication-OFF state, power spectra exhibited a prominent peak in the beta and low-gamma range (25–45 Hz; Fig. 2C). Apomorphine reduced power within this frequency range with a large effect size (1.48 ± 0.54 SD; 1.15 ± 0.41 SD; Cohen’s d = 0.66), although group-level comparisons between Med-ON and OFF conditions did not reach statistical significance. Only when the dosage was doubled in the cylinder test did beta and low-gamma suppression become significant (Supp. Fig. 1). In contrast, the therapeutic dose significantly increased high-gamma activity (0.66 ± 0.21 SD; 0.83 ± 0.19 SD; p = 0.003).

Neurobehavioral correlations revealed no significant relationship between walking distance and beta and low-gamma oscillations (Spearman r = –0.32, p = 0.093). High-gamma power, however, significantly correlated with walking distance (Spearman r = 0.50, p = 0.007). Together, these findings highlight a frequency-specific contribution of cortical oscillations to gait execution.

### Apomorphine shifts motor-state distribution towards prokinetic states

To gain deeper insight into gait-related behavioral changes, we classified locomotor behavior into three motor states: gait, akinesia, and stationary movement. States were manually annotated by a blinded rater using high-resolution video recordings (Supplementary Video 1). Apomorphine markedly shifted the distribution of time spent in each motor state (Fig. 3A). Relative to the medication-OFF condition, apomorphine increased time spent in gait (4.68 ± 2.77 SD to 11.74 ± 5.43 SD; p < 0.01), decreased stationary movement (26.39 ± 10.15 SD to 14.63 ± 10.21 SD; p < 0.01), and reduced akinesia (11.59 ± 6.53 SD to 2.41 ± 4.51 SD; p < 0.01).

**Figure 3.**
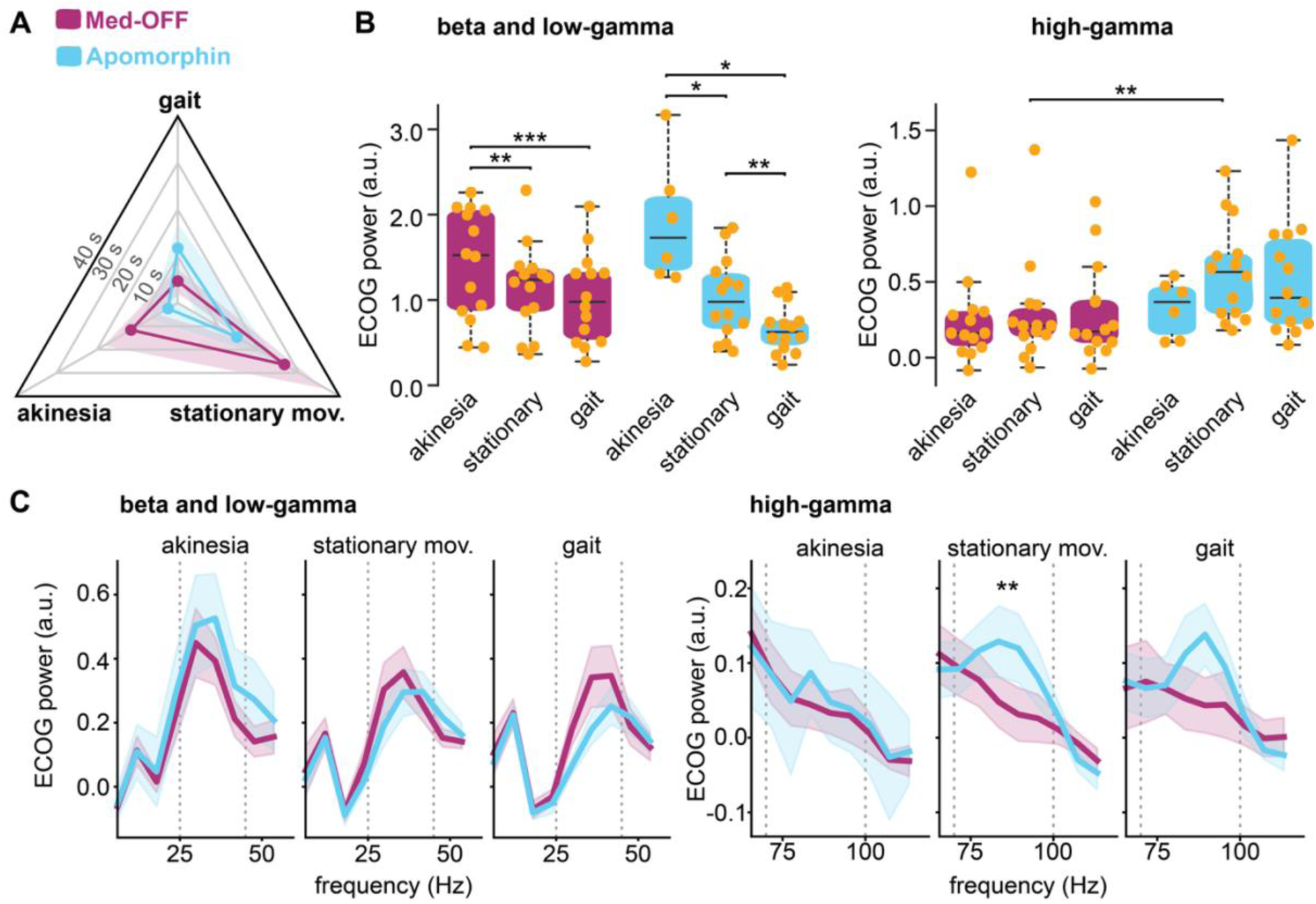
Differential modulation of cortical oscillations depends on ongoing motor states. (**A**) Apomorphine significantly altered the duration of each motor state, increasing time spent in gait and reducing stationary movement and akinesia (p < 0.01 for all pairwise comparisons). (**B**) Beta and low-gamma activity varied significantly across motor states but was not significantly affected by apomorphine. In contrast, high-gamma activity increased with apomorphine during stationary movement but did not differ significantly across motor states. (D) Power spectral densities of cortical oscillations during each motor state in Med-ON and Med-OFF conditions. Data in A and C represent mean ± SD. Significance: *p < 0.05, **p < 0.01, ***p < 0.001.

We then analyzed cortical oscillations within each motor state under Med-ON and OFF conditions(Fig. 3B and C). Two factor analysis revealed significant modulation of beta and low-gamma activity by motor state (p < 0.01), but not by apomorphine. Post hoc comparisons indicated significant differences in beta and low-gamma power between akinesia vs. stationary movement and akinesia vs. gait for both Med-OFF and ON conditions. Differences between stationary movement and gait were significant only during Med-ON recordings. High-gamma activity, in contrast, showed significant modulation by the factor drug (p < 0.001) but not by motor state. Post hoc analysis revealed that apomorphine selectively increased high-gamma power during stationary movements.

In summary, apomorphine shifted the behavioral motor-state distribution toward gait and selectively elevated high-gamma activity during stationary movement, whereas beta and low-gamma oscillations varied with motor state but were not significantly altered by the drug.

### Kinematic gait analysis reveals reduced gait velocity with apomorphine

Using 3D hindlimb kinematics, we assessed how apomorphine influenced individual gait cycle features. A total of 104 kinematic parameters (Supplementary Table 1) were extracted and subjected to principal component analysis (PCA), following previously established methods^33^. Visualization of gait patterns in PC1–PC2 space showed that PC1 explained the largest proportion of variance (26%) and significantly differentiated Med-ON from Med-OFF trials (p < 0.01; Fig. 4B). Factor loadings indicated that PC1 primarily reflected gait speed–related parameters (Fig. 4C), including gait velocity, step length, and double stance duration.

**Figure 4.**
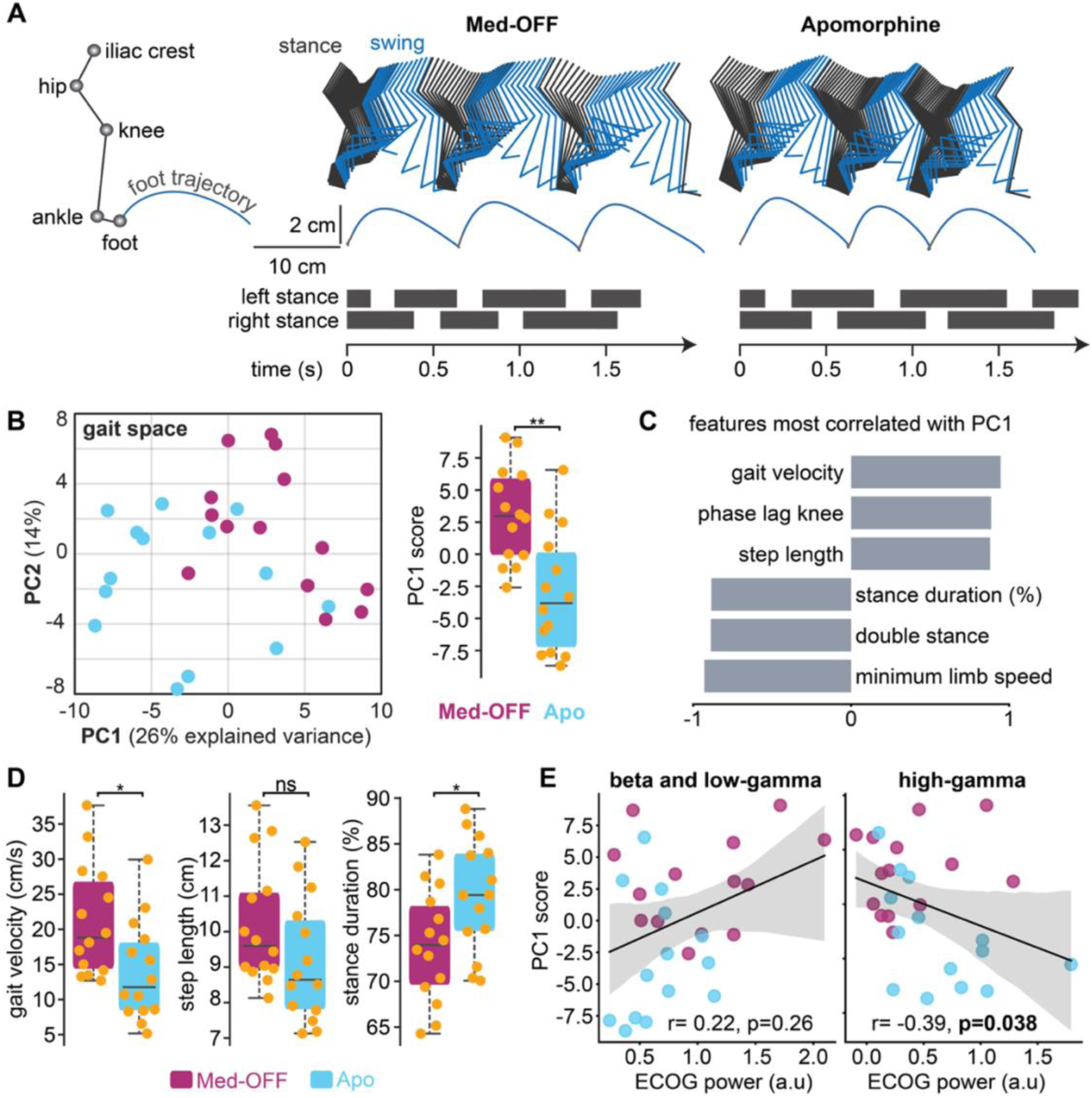
Apomorphine reduces gait speed parameters. (**A**) Left: stick figure showing hindlimb marker placement for 3D kinematic analysis. Right: representative stick sequences, foot trajectories, and gait cycle diagrams for Med-OFF and Med-ON conditions. (**B**) Principal component analysis of hindlimb kinematics revealed apomorphine-induced changes in gait pattern, primarily along PC1. (**C**) Loadings on PC1 highlighting gait-speed–related features. (E) Quantification of selected gait parameters showed significant reductions in gait velocity and increased stance duration with apomorphine; step length showed a non-significant decreasing trend. (**E**) High-gamma activity significantly correlated with PC1 scores.

Individual parameter analysis revealed that apomorphine significantly reduced gait velocity and increased stance duration, while step length was moderately reduced but did not reach significance (Fig. 4D). Correlation analyses showed no significant relationship between PC1 and beta/low-gamma activity (Spearman r = 0.22, p = 0.26; Fig. 4E), whereas high-gamma elevations significantly correlated with PC1 (Spearman r = –0.39, p = 0.038).

Together, these findings indicate that apomorphine slowed stepping behavior, and this reduction in gait speed was associated with increased high-gamma activity.

### Apomorphine altered number of motor state transitions

To further delineate the mechanisms underlying apomorphine’s effects on gait regulation, we quantified transitions among the three motor states (Fig. 5A). Apomorphine significantly reduced the total number of state transitions (Med-OFF: 5.60 ± 2.15 SD; apomorphine: 3.68 ± 1.38 SD; p = 0.004). Examining individual transition types revealed a strong reduction in transitions between stationary movement and akinesia (Med-OFF: 28.57 ± 8.49 SD; apomorphine: 6.75 ± 9.85 SD; p < 0.001), along with an increase in transitions between stationary movement and gait (Med-OFF: 2.21 ± 0.87 SD; apomorphine: 3.01 ± 0.95 SD; p < 0.05). Across the dataset, no direct transitions between gait and akinesia were observed.

**Figure 5.**
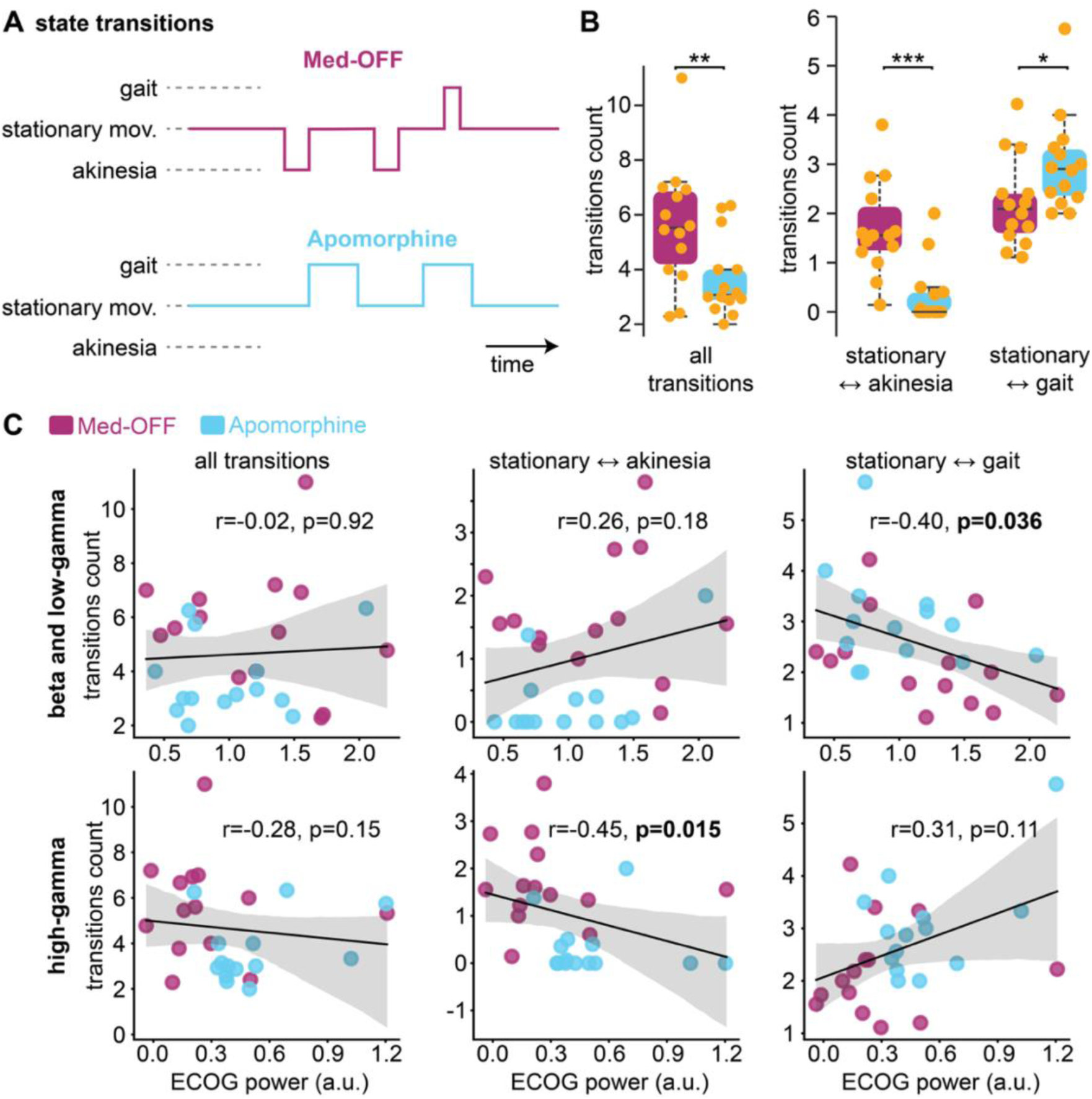
Apomorphine alters likelihood of motor state transitions. (**A**) Representative motor state sequences in Med-OFF and Med-ON conditions. (**B**) Apomorphine reduced the overall number of state transitions (left). The drug decreased transitions between stationary movement (SM) and akinesia while increasing transitions between SM and gait. (**C**) Neurobehavioral correlations showed significant negative associations between beta and low-gamma and SM→gait transitions, and between high-gamma and SM→akinesia transitions.

Correlation analyses showed that reductions in beta/low-gamma power were associated with increased transitions from stationary movement into gait (Spearman r = –0.40, p = 0.036). In contrast, elevations in high-gamma activity negatively correlated with transitions between stationary movement and akinesia (Spearman r = –0.45, p = 0.015; Fig. 5C).

Taken together, these results point to distinct roles of beta and low-gamma and high-gamma oscillations in mediating transitions between specific motor states.

## Discussion

In this study, we provide a detailed characterization of the behavioral and cortical effects of apomorphine during freely initiated locomotion in the unilateral 6-OHDA rat model of Parkinsonism. Therapeutic dosing of apomorphine increased overall walking distance on the runway, accompanied by a redistribution of time spent across motor behaviors with more time in gait and less time in stationary movement and akinesia. Kinematic analysis further showed that, despite enhancing the occurrence of gait, apomorphine reduced gait speed. At the neural level, cortical recordings revealed a robust increase in high-gamma activity with apomorphine, whereas beta and low-gamma oscillations remained largely unchanged by the drug but varied across motor states. Correlational analyses linked reductions in beta and low-gamma activity with the initiation of gait and elevations in high-gamma activity with reduced transitions into akinesia. Collectively, the data reveal a complex pattern of apomorphine-induced changes spanning motor states, kinematics, and cortical oscillations. We reason that these effects are best interpreted as a shift in the underlying motor-state equilibrium.

### Concept of a motor-state equilibrium for gait regulation in Parkinson’s disease

Previous literature supports the status quo hypothesis, which posits that beta oscillations stabilize the current motor state in Parkinson’s disease^14^. In the akinetic OFF state, cortical - basal ganglia coherence is dominated by beta-band activity^1,13,14^, whereas levodopa reduces beta power and introduces a gamma-band peak associated with movement facilitation^18,34^. Our findings in the unilateral 6-OHDA model extend this framework to self-initiated locomotion. In this paradigm, we identified three principal motor states - akinesia, stationary movement, and gait - that were separable at both behavioral and neural levels. Rather than characterizing oscillatory bands as uniformly prokinetic or antikinetic, we reason that cortical oscillations regulate locomotion in a state specific manner. To better illustrate this idea, we propose the concept of a motor-state equilibrium that is defined by the durations of and transitions between states (Fig. 6). Within this framework, the behavioral effects of apomorphine can be interpreted as a drug-induced shift in this equilibrium driven by state-specific oscillatory patterns. High-gamma activity, for example, was selectively elevated during stationary movement and suppressed transitions into akinesia, yet, it did not significantly contribute to transitions into gait. In contrast, gait initiation was associated with proportional reduction in beta and low-gamma activity. Although apomorphine reduced akinesia, this effect did not depend on a drug-induced suppression of beta and low-gamma power during the akinetic state. Together, these findings underscore the value of a state-specific perspective for understanding cortical activity and gait-related therapies.

**Figure 6.**
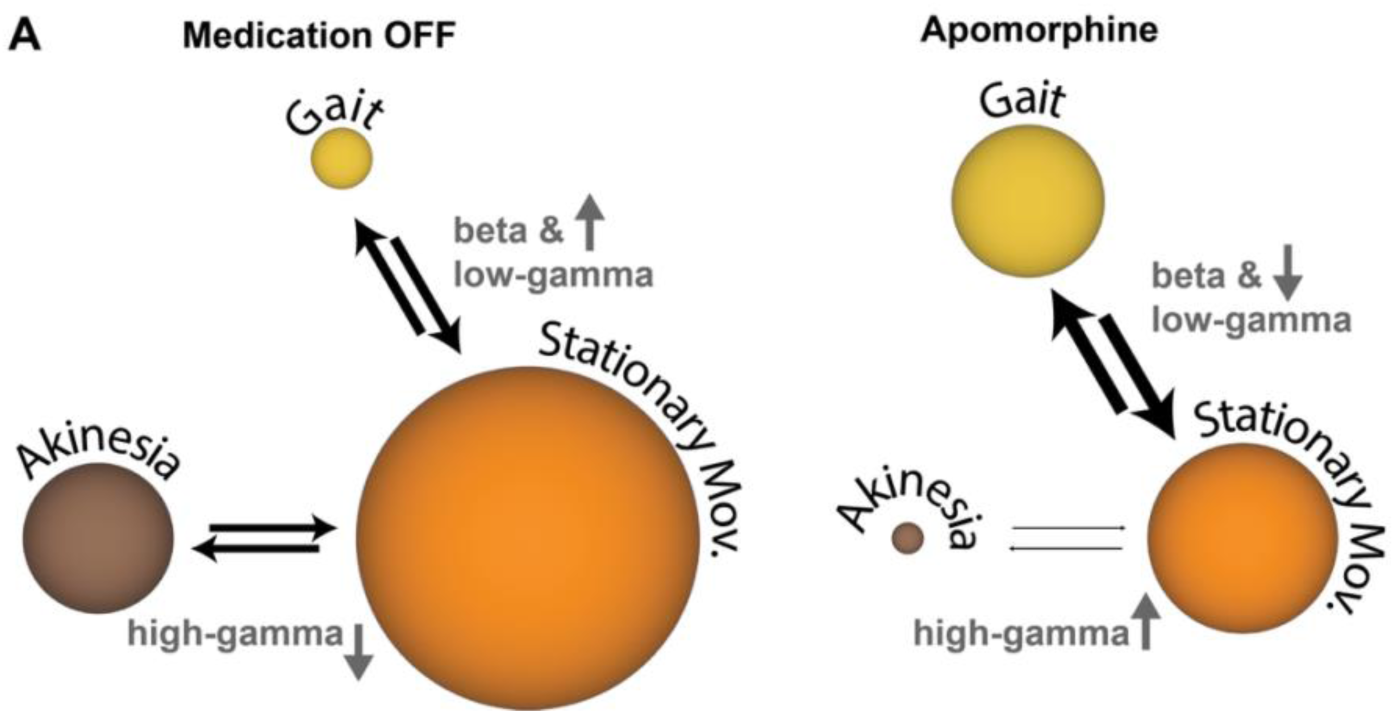
Modulation of motor state equilibrium by apomorphine. (**A**) Schematic representation of motor state equilibrium defined by the observed states, their durations, and state-transition likelihoods. Within this framework, cortical oscillations regulate the probability of transitioning between states, thereby extending or reducing the time spent in each motor state. Colors indicate motor states; sphere size represents average state duration; arrow width reflects transition likelihood.

### Network components involved in apomorphine gait effects

Our observations showed that apomorphine promoted gait initiation, prolonged gait episodes, yet, simultaneously slowed gait at the kinematic level. These behavioral effects may, in part, reflect the distinct contributions of D1-and D2-type dopaminergic receptors within cortico - basal ganglia loops^35^. Activation of striatal D1 receptors facilitates locomotor initiation^36^, and both striatal D1 and D2 receptors contribute to the maintenance of gait^35^. Given the receptor affinities of apomorphine, it is likely that both receptor classes contributed to the effects observed in our study, with a predominance for D2 activation. In principle, apomorphine may also act directly on cortical dopamine receptors. Activation of either cortical D1 or D2 receptors increases the excitability of layer 5 pyramidal neurons^37^. However, cortical dopamine signaling has been linked primarily to motor learning and fine motor control ^37^, and its specific role in regulating gait remains to be investigated. Together, cortico–basal ganglia mechanisms provide a plausible explanation for apomorphine’s effects on motor state regulation.

A second region of conceptual relevance is the spinal cord, which expresses both D1 and D2 receptors and receives its primary dopaminergic input from the diencephalic Area 11^38^. Optogenetic activation of this region initiates locomotion - an effect attributed mainly to Gs-coupled D1 receptor in the spinal cord^38,39^. In contrast, spinal D2 receptors have been shown to slow locomotor output^40^, offering a potential mechanism for the kinematic slowing we observed with apomorphine.

In summary, it is plausible that apomorphine modulates multiple components of the locomotor network in distinct and potentially opposing ways. Whether these divergent effects on motor state regulation and gait kinematics translate to human patients - and whether they ultimately yield a net therapeutic benefit for gait - remains an important question for future work.

### Towards state-adaptive neuromodulation strategies in Parkinson’s disease

Adaptive deep brain stimulation (aDBS) has demonstrated improved motor outcomes in Parkinson’s disease compared to standard, continuous stimulation^41,42^. Most aDBS approaches rely on monitoring a specific biomarker - often beta-band power - and triggering stimulation when the signal exceeds a predefined threshold^41^. Refining this approach by tuning stimulation to features of beta bursts, including their amplitude and duration, has shown clinical benefits for freezing of gait, with efficacy comparable to continuous DBS^43^. More recent concepts aim to broaden adaptive neuromodulation by incorporating symptom-specific frequency bands, as well as contextual variables such as behavioral activities, symptom fluctuations, and sleep–wake cycles^44,45^. Our data suggest that beta-and gamma-band activity exert state-dependent influences during locomotion, differentially regulating transitions into akinesia or gait. Although these observations arise from a pharmacological manipulation, they raise the question of whether DBS can elicit comparable state-dependent modulation of brain activity during gait. Exploiting state-specific frequency signatures may offer a path toward next-generation adaptive neuromodulation strategies that more precisely target gait control in Parkinson’s disease.

### Limitations of the study

Our study shows that apomorphine robustly modulates high-gamma activity but does not significantly alter beta or low-gamma activity at the therapeutic dose used. This pattern should be viewed as dose-dependent. Prior literature^7^ and our own data indicate that higher doses of apomorphine can suppress beta and low-gamma power (see Supplementary Fig. 1). However, such suppression is often accompanied by unilateral body rotations, a behavior that complicates the interpretation of dopaminergic treatment effects in the Parkinsonian phenotype. For this reason, higher doses are less suitable for investigating gait-therapy mechanisms in the 6-OHDA model.

Another limitation relates to the translational comparability of locomotor deficits in rodent models and human Parkinson’s disease. While bradykinesia during gait is readily quantifiable in rodents, comparability for freezing of gait (FoG) is less direct. In patients, FoG involves a sudden interruption of ongoing gait - manifesting as trembling, shuffling, festination, or an akinetic episode^3^. In our rodent model, we observed akinetic episodes, but these did not occur as abrupt interruptions of gait; instead, they emerged through transitions into stationary movement. Thus, rather than modeling FoG per se, our findings may better translate to more general disruptions of the sequence and continuity of gait.

## Conclusion and translational outlook

Our study demonstrates that the gait-therapy effects of apomorphine are best understood as a shift in the motor-state equilibrium, with cortical oscillations exerting state-dependent influences on locomotor regulation. These findings outline a mechanistic framework for understanding gait modulation in Parkinson’s disease and suggest new avenues for the development of targeted, state-informed gait therapies.

## Acknowledgments

N.W., C.H., and M.E. conceived and designed the study. B.K. performed the lesion surgeries and electrode implantations. B.K. and E.L.G. acquired data. B.K., E.L.G., and M.G. analyzed data. R.D.S., A.V., R.B., M.S., and R.P. provided technical support. N.W., C.H., and M.E. secured funding and resources. N.W. supervised the project. B.K. and N.W. wrote the manuscript. All authors edited and approved the final version of the manuscript.

## Sources of funding

E.L.G., B.K., W.J.N., M.E., C.H., and N.W. received support from the Collaborative Research Center ReTune (TRR 295, project number 424778381). N.W. received support form the Volkswagen-Foundation (Freigeist Fellowship) and the tenure-track program (German Federal Ministry of Education and Research, BMBF). W.J.N. was additionally funded by the European Research Council (ERC, ReinforceBG, project 101077060) and BMBF (project FKZ01GQ1802). M.E. received further support from the German Research Foundation (DFG) under Germany’s Excellence Strategy (EXC-2049 – 390688087) and the Clinical Research Group KFO 5023 BeCAUSE-Y (project 2, EN343/16-1), as well as from the BMBF, DZNE, DZHK, DZPG, the EU, the Corona Foundation, and Fondation Leducq. C.H. was funded by the DFG (Excellence Strategy, EXC-2049 – 390688087, SPARK-BIH Stroke Protect), the EraNet Neuron project (DFG project 522473931), the DZHK/BMBF Innovation Cluster (FKZ81X2100285), Charité 3R (C3RHeaD and STROKE-PREDICT-C3R), the BIH, and the Fondation Leducq Transatlantic Network of Excellence (17CVD03).

## Conflict of interest

W.J.N. received honoraria for consulting from InBrain–Neuroelectronics that is a neurotechnology company and honoraria for talks from Medtronic that is a manufacturer of deep brain stimulation devices, both unrelated to this manuscript. N.W. is co-inventor of two patents on spinal cord stimulation technologies for gait therapy.

## Declaration of generative AI and AI-assisted technologies in the writing process

During the preparation of this work the author(s) used OpenAI / ChatGPT-4o to improve language and readability. After using this tool, the authors further reviewed and edited the manuscript and figures and take full responsibility for the content of the publication.

## Supplementary Information

**Supplementary Figure 1.**
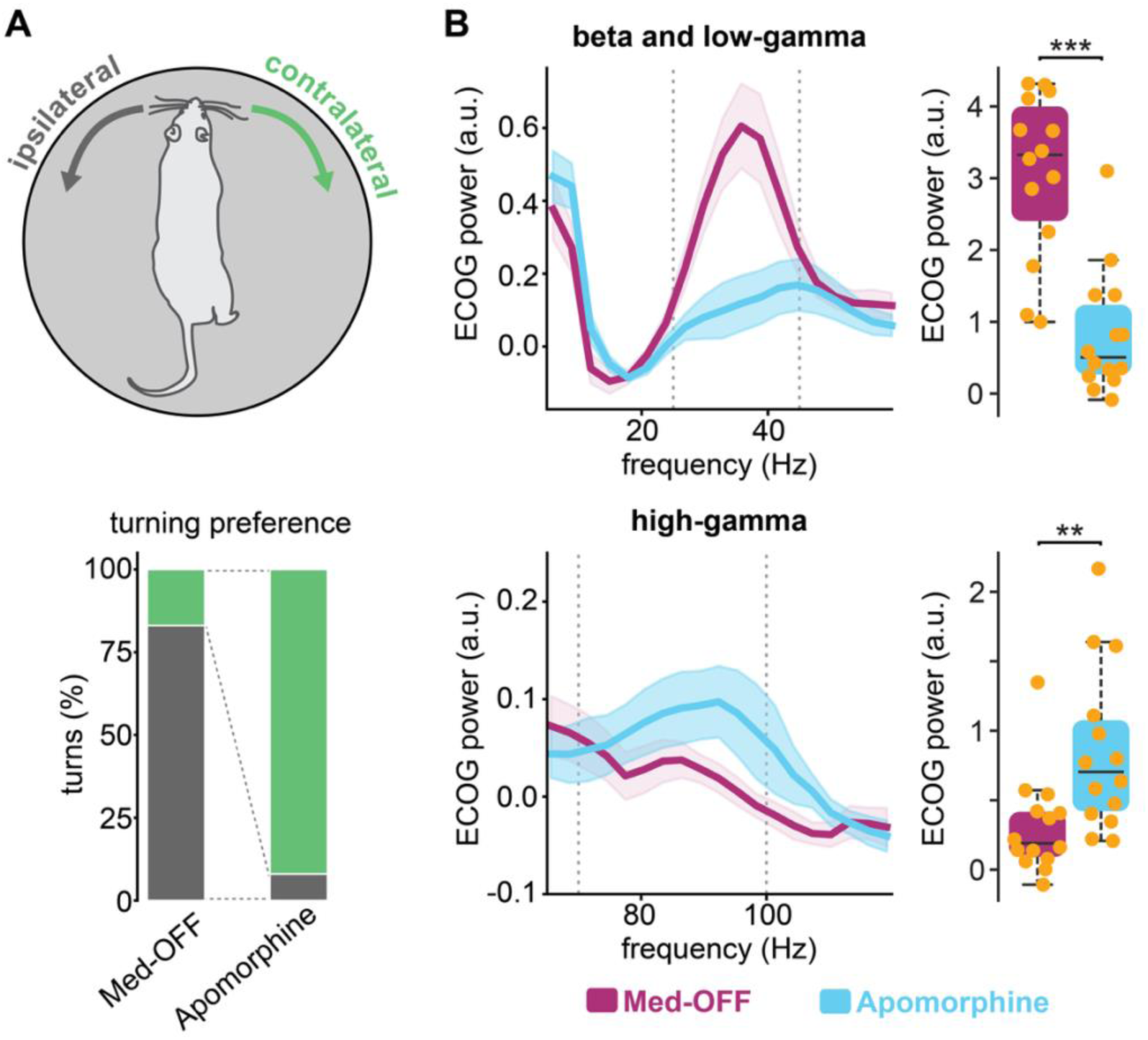
Cylinder test. (**A**) Top: schematic of a rat inside the cylinder. Gray arrow indicates ipsilateral turns (toward the lesioned hemisphere); green arrow indicates contralateral turns. Bottom: proportion of ipsilateral versus contralateral turns under both conditions. (**B**) Left: power spectral density (PSD) plots for beta and low-gamma (HB-LG) and high-gamma (HG) ranges during 5-minute recordings. Gray dashed lines mark 25 and 45 Hz (HB-LG) and 70 and 100 Hz (HG). Right: quantification of HB-LG and HG activity as area under the curve. Magenta shows baseline, blue shows apomorphine. Individual datapoints shown in light orange. Boxplots show median and interquartile ranges. Significance: **p < 0.01, ***p < 0.001.

**Supplementary Video 1.** Example of a unilateral 6-OHDA rat walking across the runway under medication OFF and apomorphine conditions. The behavioral changes described in the study can be observed, including apomorphine-induced reductions in akinesia and prolonged gait episodes, consistent with a shift in the motor-state equilibrium. (Video link: https://youtu.be/x48_jwCESz4)

---

**Supplementary Table 1.**
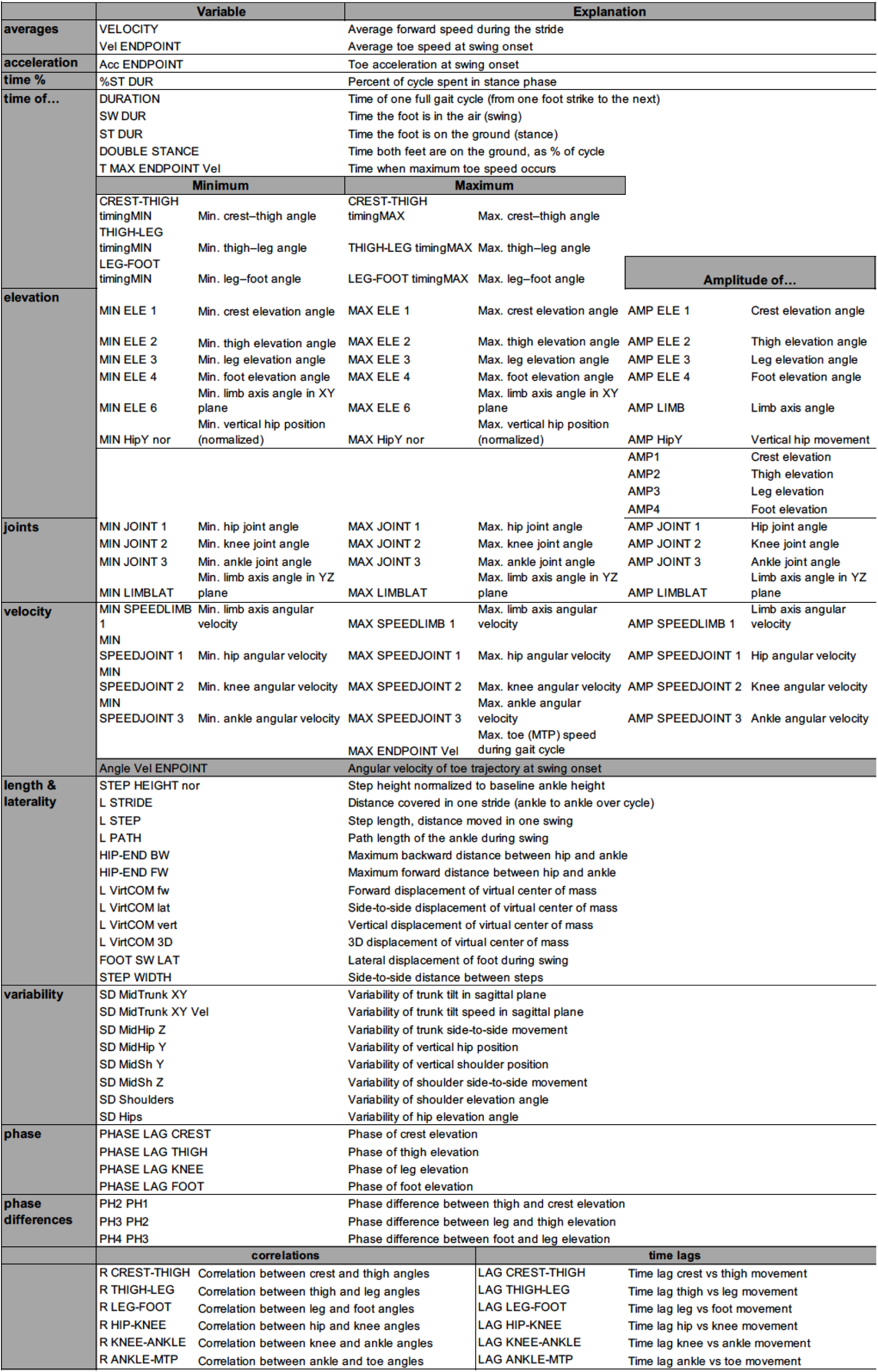
List of all parameters for kinematic gait analysis.

## Notes

### Competing Interest Statement

The authors have declared no competing interest.

### Summary of Updates

Discussion revised; supplemental files updates; authors added.

## Citations

1. Moustafa AA, Chakravarthy S, Phillips JR, Gupta A, Keri S, Polner B, Frank MJ, Jahanshahi M. Motor symptoms in Parkinson’s disease: A unified framework. Neuroscience & Biobehavioral Reviews. 2016;68:727–740.

2. Hardeman LES, Kal EC, Young WR, van der Kamp J, Ellmers TJ. Visuomotor control of walking in Parkinson’s disease: Exploring possible links between conscious movement processing and freezing of gait. Behavioural Brain Research. 2020;395:112837.

3. Nutt JG, Bloem BR, Giladi N, Hallett M, Horak FB, Nieuwboer A. Freezing of gait: moving forward on a mysterious clinical phenomenon. Lancet Neurol. 2011;10:734–744.

4. Pozzi NG, Canessa A, Palmisano C, Brumberg J, Steigerwald F, Reich MM, Minafra B, Pacchetti C, Pezzoli G, Volkmann J, et al. Freezing of gait in Parkinson’s disease reflects a sudden derangement of locomotor network dynamics. Brain. 2019;142:2037–2050.

5. Mirelman A, Bonato P, Camicioli R, Ellis TD, Giladi N, Hamilton JL, Hass CJ, Hausdorff JM, Pelosin E, Almeida QJ. Gait impairments in Parkinson’s disease. Lancet Neurol. 2019;18:697–708.

6. Pötter-Nerger M, Volkmann J. Deep brain stimulation for gait and postural symptoms in Parkinson’s disease. Mov Disord. 2013;28:1609–1615.

7. Wenger N, Vogt A, Skrobot M, Garulli EL, Kabaoglu B, Salchow-Hömmen C, Schauer T, Kroneberg D, Schuhmann MK, Ip CW, et al. Rodent models for gait network disorders in Parkinson’s disease – a translational perspective. Experimental Neurology. 2022;352:114011.

8. Auffret M. Apomorphine: past, present and future in Parkinson’s disease and beyond. Parkinsonism & Related Disorders. 2025;139:107872.

9. Jenner P, Katzenschlager R. Apomorphine - pharmacological properties and clinical trials in Parkinson’s disease. Parkinsonism & Related Disorders. 2016;33:S13–S21.

10. Katzenschlager R, Poewe W, Rascol O, Trenkwalder C, Deuschl G, Chaudhuri KR, Henriksen T, Van Laar T, Spivey K, Vel S, et al. Apomorphine subcutaneous infusion in patients with Parkinson’s disease with persistent motor fluctuations (TOLEDO): a multicentre, double-blind, randomised, placebo-controlled trial. The Lancet Neurology. 2018;17:749–759.

11. Isaacson SH, Espay AJ, Pahwa R, Agarwal P, Shill HA, Hui J, Dashtipour K, Lew M, Qin P, Formella AE, et al. Continuous, subcutaneous apomorphine infusion for Parkinson disease motor fluctuations: Results from the phase 3, long-term, open-label United States InfusON study. Journal of Parkinson’s Disease. 2025;15:361–373.

12. Corboy DL, Wagner ML, Sage JI. Apomorphine for Motor Fluctuations and Freezing in Parkinson’s Disease. Ann Pharmacother. 1995;29:282–288.

13. Kühn AA, Kupsch A, Schneider G-H, Brown P. Reduction in subthalamic 8-35 Hz oscillatory activity correlates with clinical improvement in Parkinson’s disease. Eur J Neurosci. 2006;23:1956–1960.

14. Brown P, Oliviero A, Mazzone P, Insola A, Tonali P, Di Lazzaro V. Dopamine dependency of oscillations between subthalamic nucleus and pallidum in Parkinson’s disease. J Neurosci. 2001;21:1033–1038.

15. Mathiopoulou V, Lofredi R, Feldmann LK, Habets J, Darcy N, Neumann W-J, Faust K, Schneider G-H, Kühn AA. Modulation of subthalamic beta oscillations by movement, dopamine, and deep brain stimulation in Parkinson’s disease. npj Parkinson’s Disease. 2024;10:77.

16. Pogosyan A, Gaynor LD, Eusebio A, Brown P. Boosting cortical activity at Beta-band frequencies slows movement in humans. Curr Biol. 2009;19:1637–1641.

17. Engel AK, Fries P. Beta-band oscillations--signalling the status quo? Curr Opin Neurobiol. 2010;20:156–165.

18. Muthuraman M, Bange M, Koirala N, Ciolac D, Pintea B, Glaser M, Tinkhauser G, Brown P, Deuschl G, Groppa S. Cross-frequency coupling between gamma oscillations and deep brain stimulation frequency in Parkinson’s disease. Brain. 2020;143:3393–3407.

19. Acharyya P, Daley KW, Choi JW, Wilkins KB, Karjagi S, Cui C, Seo G, Abay AK, Bronte-Stewart HM. Closing the loop in DBS: A data-driven approach. Parkinsonism & Related Disorders. 2025;134:107348.

20. Thenaisie Y, Lee K, Moerman C, Scafa S, Gálvez A, Pirondini E, Burri M, Ravier J, Puiatti A, Accolla E, et al. Principles of gait encoding in the subthalamic nucleus of people with Parkinson’s disease. Sci. Transl. Med. 2022;14:eabo1800.

21. Yin Z, Zhu G, Liu Y, Zhao B, Liu D, Bai Y, Zhang Q, Shi L, Feng T, Yang A, et al. Cortical phase-amplitude coupling is key to the occurrence and treatment of freezing of gait. Brain. 2022;145:2407–2421.

22. Jiang X, Yang J, Wang Z, Jia J, Wang G. Functional interaction of abnormal beta and gamma oscillations on bradykinesia in parkinsonian rats. Brain Res Bull. 2024;209:110911.

23. Dupre KB, Cruz AV, McCoy AJ, Delaville C, Gerber CM, Eyring KW, Walters JR. Effects of L-dopa priming on cortical high beta and high gamma oscillatory activity in a rodent model of Parkinson’s disease. Neurobiol Dis. 2016;86:1–15.

24. Brazhnik E, Novikov N, McCoy AJ, Ilieva NM, Ghraib MW, Walters JR. Early decreases in cortical mid-gamma peaks coincide with the onset of motor deficits and precede exaggerated beta build-up in rat models for Parkinson’s disease. Neurobiology of Disease. 2021;155:105393.

25. Lemaire N, Hernandez LF, Hu D, Kubota Y, Howe MW, Graybiel AM. Effects of dopamine depletion on LFP oscillations in striatum are task-and learning-dependent and selectively reversed by ¡span class=“smallcaps smallerCapital”¿l¡/span¿-DOPA. Proceedings of the National Academy of Sciences. 2012;109:18126–18131.

26. Li Q, Ke Y, Chan DCW, Qian Z-M, Yung KKL, Ko H, Arbuthnott GW, Yung W-H. Therapeutic deep brain stimulation in Parkinsonian rats directly influences motor cortex. Neuron. 2012;76:1030–1041.

27. Delaville C, McCoy AJ, Gerber CM, Cruz AV, Walters JR. Subthalamic Nucleus Activity in the Awake Hemiparkinsonian Rat: Relationships with Motor and Cognitive Networks. J. Neurosci. 2015;35:6918–6930.

28. Güttler C, Altschüler J, Tanev K, Böckmann S, Haumesser JK, Nikulin VV, Kühn AA, Riesen C van. Levodopa-Induced Dyskinesia Are Mediated by Cortical Gamma Oscillations in Experimental Parkinsonism. Movement Disorders. 2021;36:927–937.

29. Skovgård K, Barrientos SA, Petersson P, Halje P, Cenci MA. Distinctive Effects of D1 and D2 Receptor Agonists on Cortico-Basal Ganglia Oscillations in a Rodent Model of L-DOPA-Induced Dyskinesia. Neurotherapeutics. 2023;20:304–324.

30. van den Brand R, Heutschi J, Barraud Q, DiGiovanna J, Bartholdi K, Huerlimann M, Friedli L, Vollenweider I, Moraud EM, Duis S, et al. Restoring voluntary control of locomotion after paralyzing spinal cord injury. Science. 2012;336:1182–1185.

31. Mathis A, Mamidanna P, Cury KM, Abe T, Murthy VN, Mathis MW, Bethge M. DeepLabCut: markerless pose estimation of user-defined body parts with deep learning. Nat Neurosci. 2018;21:1281–1289.

32. Donoghue T, Haller M, Peterson EJ, Varma P, Sebastian P, Gao R, Noto T, Lara AH, Wallis JD, Knight RT, et al. Parameterizing neural power spectra into periodic and aperiodic components. Nature Neuroscience. 2020;23:1655–1665.

33. Wenger N, Moraud EM, Raspopovic S, Bonizzato M, DiGiovanna J, Musienko P, Morari M, Micera S, Courtine G. Closed-loop neuromodulation of spinal sensorimotor circuits controls refined locomotion after complete spinal cord injury. Science Translational Medicine. 2014;6:255ra133.

34. Lofredi R, Neumann W-J, Bock A, Horn A, Huebl J, Siegert S, Schneider G-H, Krauss JK, Kühn AA. Dopamine-dependent scaling of subthalamic gamma bursts with movement velocity in patients with Parkinson’s disease. eLife. 2018;7:e31895.

35. Zhang H, Su Y, Wu X, Gan W-B. The regulation of rhythmic locomotion by motor cortical and dopaminergic inputs in the mouse striatum. Mol Brain. 2025;18:63.

36. Kravitz AV, Freeze BS, Parker PRL, Kay K, Thwin MT, Deisseroth K, Kreitzer AC. Regulation of parkinsonian motor behaviours by optogenetic control of basal ganglia circuitry. Nature. 2010;466:622–626.

37. Cousineau J, Plateau V, Baufreton J, Le Bon-Jégo M. Dopaminergic modulation of primary motor cortex: From cellular and synaptic mechanisms underlying motor learning to cognitive symptoms in Parkinson’s disease. Neurobiology of Disease. 2022;167:105674.

38. Koblinger K, Jean-Xavier C, Sharma S, Füzesi T, Young L, Eaton SEA, Kwok CHT, Bains JS, Whelan PJ. Optogenetic Activation of A11 Region Increases Motor Activity. Frontiers in Neural Circuits [Internet]. 2018;12. Available from: https://www.frontiersin.org/articles/10.3389/fncir.2018.00086

39. Han P, Nakanishi ST, Tran MA, Whelan PJ. Dopaminergic Modulation of Spinal Neuronal Excitability. J. Neurosci. 2007;27:13192–13204.

40. Sharples SA, Humphreys JM, Jensen AM, Dhoopar S, Delaloye N, Clemens S, Whelan PJ. Dopaminergic modulation of locomotor network activity in the neonatal mouse spinal cord. Journal of Neurophysiology. 2015;113:2500–2510.

41. Little S, Pogosyan A, Neal S, Zavala B, Zrinzo L, Hariz M, Foltynie T, Limousin P, Ashkan K, FitzGerald J, et al. Adaptive deep brain stimulation in advanced Parkinson disease. Ann Neurol. 2013;74:449–457.

42. Acharyya P, Daley KW, Choi JW, Wilkins KB, Karjagi S, Cui C, Seo G, Abay AK, Bronte-Stewart HM. Closing the loop in DBS: A data-driven approach. Parkinsonism & Related Disorders. 2025;134:107348.

43. Wilkins KB, Petrucci MN, Lambert EF, Melbourne JA, Gala AS, Akella P, Parisi L, Cui C, Kehnemouyi YM, Hoffman SL, et al. Beta burst-driven adaptive deep brain stimulation for gait impairment and freezing of gait in Parkinson’s disease. Brain Communications. 2025;7:fcaf266.

44. Neumann W, Gilron R, Little S, Tinkhauser G. Adaptive Deep Brain Stimulation: From Experimental Evidence Toward Practical Implementation. Movement Disorders. 2023;38:937–948.

45. Tinkhauser G, Moraud EM. Controlling Clinical States Governed by Different Temporal Dynamics With Closed-Loop Deep Brain Stimulation: A Principled Framework. Front. Neurosci. 2021;15:734186.

